# Gut Microbiota Succession and Metabolic Pathway Remodeling in the Progression of Type 2 Diabetes Mellitus in Adult Rats

**DOI:** 10.1101/2025.07.08.663737

**Authors:** Yanchao Liu, Han Bao, Jinni Yao, Yang Jiao, Hailing Li, Tao Yan

## Abstract

**Background:** Emerging evidence highlights the complex interplay between gut microbiota and type 2 diabetes mellitus (T2D); however, the temporal trajectory of microbial community succession and the concomitant shifts in associated metabolic pathways across the disease lifecycle remain poorly characterized. In this study, we conducted a multi-dimensional analysis of gut microbiota composition and functional pathways in rats, spanning the continuum from health, through the T2D pathological state, to post-intervention recovery phases.

**Results:** Healthy aged rats exhibited significantly greater gut microbial diversity compared to both younger healthy cohorts and T2D rats. In the absence of intervention, progressive T2D development led to a sustained decline in microbial diversity that persisted until death. Notably, β-tocotrienol (β-T3), metformin hydrochloride (MET) and healthy dietary interventions not only improved fasting blood glucose (FPG) levels in T2D mice but also restored their gut microbial diversity and metabolic functions. At the phylum level, the ratio of p Bacteroidota to p Firmicutes exhibited an age-dependent increase in healthy rats but displayed a progressive decline in T2D rats. Notably, species-specific expansions were observed across both healthy and pathological states. For example, *s Bacteroides_acidifaciens* (*B. acidifaciens*) showed a significant expansion in early T2D stage disease, while *s Anaerostipes_hadrus* became the dominant species in advanced-stage T2D. Among intervention groups, *s Phascolarctobacterium_faecium* was enriched in the β-tocotrienol (β-T3) group; the abundance of *s Escherichia_coli* (*E. coli*) increased dramatically in the deceased rats. Gut metabolites also exhibited distinct characteristics across different physiological stages. In young rats, the metabolite profile was characterized by lower abundances of most metabolites, which stabilized with age accompanied by upregulation of select molecules in older rats. Notably, during T2D progression, the metabolite profile displayed a largely opposite pattern compared to healthy controls. Importantly, dietary interventions including healthy food, β-T3, MET partially restored metabolic homeostasis in T2D rats. Further analysis revealed that key differential metabolites across groups were enriched in bile acids (BAs), amino acids, benzene and its substituted derivatives, and organic acids with their derivatives. Specifically, amino acid metabolism was more active in healthy rats, whereas BA and benzene derivative metabolism were upregulated in T2D. Key microbial species driving these shifts included *s Lactobacillus_johnsonii*, *s Lactobacillus_reuteri,* and *B. acidifaciens*, which exhibited positive correlations with BAs and select amino acids. The *B. acidifaciens* could be a biomarker of T2D and glycoursodeoxycholic acid, sohyodeoxycholic acid, beta-muricholic acid and allocholic acid were the key BAs to regulate glycometabolism of the host.

**Conclusions:** During the T2D stage, both gut microbiota composition and metabolic profiles exhibited distinct characteristics compared to those of healthy age-matched older rats. Notably, β-T3 and MET interventions not only facilitated the recovery of gut microbial diversity and metabolite levels in T2D mice but also promoted the proliferation of specific dominant species, each intervention favoring distinct taxa. In this context, *B. acidifaciens* emerged as a dominant bacterial species in T2D, exerting regulatory effects on glycometabolism through BA and amino acid metabolic pathways. Collectively, these findings advance our understanding of T2D pathophysiology and highlight the potential of probiotic-based therapies for its management.

**Importance:** The T2D is a kind of chronic disease and the gut microbiota shows a continuously slow succession from some microbial populations to other ones in the disease progress. Uncovering the rule of change in gut bacteria from healthy rats to T2D ones and the recovering capacity of gut microbiota through interventions of healthy food, β-T3 and antidiabetic drug are beneficial to understand the real regulation mechanism of gut bacteria in T2D progress. Then the results lay the foundation of developing probiotics and researching new metabolic therapeutic target of gut microorganisms. In addition, to research the metabolic regulation mechanism of gut bacteria, using multi-omics including microbiomics and metabonomics is an effective way.

## Introduction

T2D is a main kind of diabetes (over 90%) with symptom of hyperglycaemia, and 1 in 10 people are living with diabetes worldwide (1). The prevalence of diabetes is increasing in the coming years, and the patients will be 783 million in 2045 (1). To control the serious disease, many researches focus on the new targets of prevention and therapy (2, 3). Undoubtedly, gut microbiota has been a hot point(4). In T2D patients, the gut microbiota is slight dysbacteriosis, number of beneficial bacteria decrease like butyrate-producing bacteria (5). There are several hypotheses to explain the mechanisms of gut microbiota regulating the T2D. One is gut microorganisms-derived metabolites, like short-chain fatty acids (SCFA) and bile acids (BAs) that play a role of communication medium between the gut bacteria and the hosts (6, 7). SCFAs mainly come from metabolism of intestinal anaerobes, and the butyrate can protect islet cells from damage (8). BAs can regulate lipid and glucose metabolism by activating the gut farnesol X receptor (FXR) and G protein-coupled bile acid receptor 1 (9). The other is lipopolysaccharide (LPS), that can disturb the gut barrier by inducing mild inflammatory reaction (6, 7). Although the relationship between gut bacteria and T2D was proved by lots of researches, the rules of gut microbial communities succession in the process of T2D is unclear. In addition, to understand the forces that shape the gut microbiome, especially at the strain level is an imperative (10). And the metabolic mechanisms of some specific gut microbial species regulating glycometabolism also need more studies to uncover. Except SCFAs and BAs, there also other metabolites in gut could be related with glycometabolism of the host (11). Our previous research showed that intestinal bacteria-derived β-tocotrienol (β-T3) and its metabolic pathway had correlation with T2D (12).

## Methods

### Animal experiment

Experimental rats and fodders were all bought from SPF (Beijing) Biotechnology Co.,Ltd.. Two months (8 weeks) old Wistar mouses were acclimatized under conditions at 20-25℃, 50% relative humidity, and 12/12 h light/dark cycle for 1 week(13), then the rats were randomly divided into BL (base line group), CT (control group, 17 weeks), CF (control group, 21 weeks) and model groups, using random number table. BL, CT and CF were fed by maintenance fodder (MF) for 0, 8, and 12 weeks respectively, each group contained 6 rats. The rats of BL were killed to collect colons after adaptive feeding for a week. The model groups were used to induce T2D rats, 36 rats totally, fed by T2D model fodder SFD019 (TMF, MF 67%, Lard 10%, Sucrose 20%, cholesterol 2.5%, Sodium cholate 0.5%) for 8 weeks, then injected streptozotocin (STZ, January 30, 2024, 30mg/kg) into each enterocoelia of rat. FPG values were tested using caudal tip blood, after fasting for 12 hours. When the FPG upped at least 16.7mmol/L, 6 rats were randomly selected to be T2D group and killed to get their colons. Meanwhile, the CT also was treated similarly. The remainder we divided them into 4 groups, (1) RFI: MF instead of TMF, 6 rats; (2) bT3I:

MF and β-T3 (0.25mg dissolving in 1ml corn oil and feeding that to each rat by gavage every day), 6 rats; (3) MI: MF and MET (45mg each mouse each day) 6 rats; (4) HFI: continually fed TMF, 12 rats. The four groups all intervened for 3 weeks. At the end, only 3 still lived from the HFI (12 rats), they were still named HFI, and the other died ones were random selected 6 named the Death group. The CF, RFI, bT3I, MI, HFI and Death 6 groups were killed to collect colons at the same time, in the end of the experiment. All the BL, CT, T2D, CF, RFI, bT3I, MI, HFI and Death 9 groups rats were anesthetized by injecting isoflurane (0.3ml/100g) and collected feces squeezed from colons to test microbiota and metabolites. The experiment was started December 6, 2023, and ended February 28, 2024, for 12 weeks. At every weekend we measured both FPG value and weight of each mouse (figure 1).

**Figure 1.**
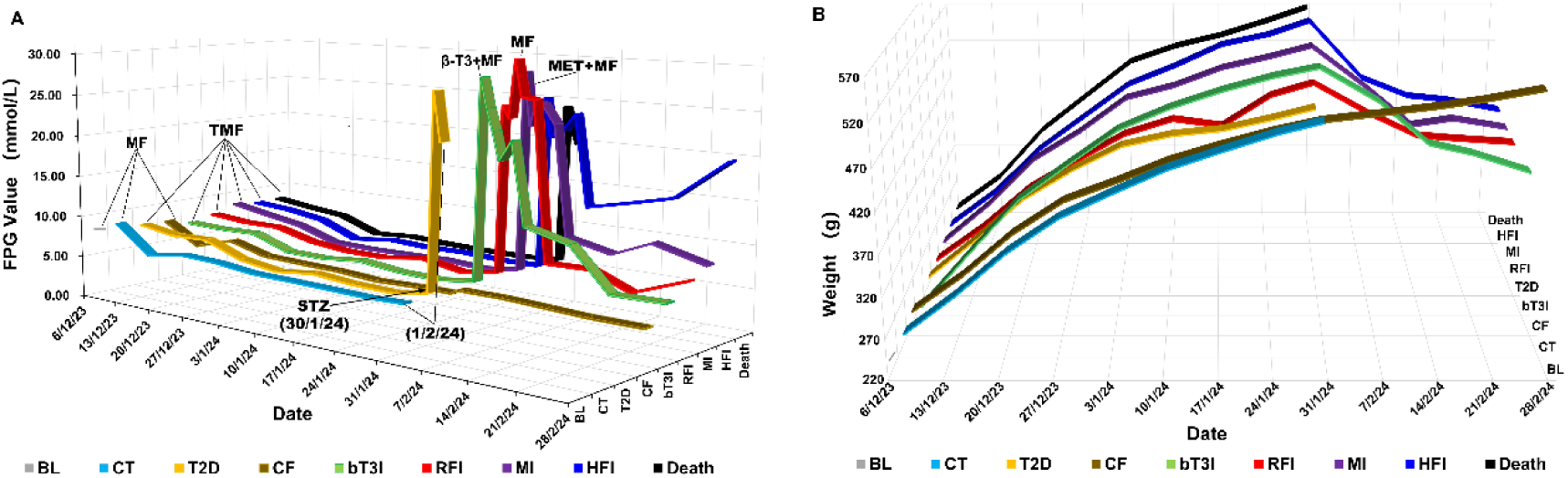
Temporal changes in fasting plasma glucose (FPG) levels and body weights of each group at key time points during the experimental progression (9 groups total). FPG and body weight measurements were collected at the same time points across groups. MF: Maintenance diet feeding; TMF: T2D model diet feeding; MET + MF: Co-administration of maintenance diet and MET; β-T3 + MF: Concurrent administration of maintenance diet and β-T3. STZ was administered on January 30, 2024, and successful T2D induction in rats was confirmed by February 5, 2024 (1 week post-injection). Interventions (MET or β-T3 treatment) were initiated on February 5, 2024, and continued for 3 weeks until February 28, 2024. A. Temporal dynamics of FPG levels over the entire experimental period. B. Temporal changes in body weight over the entire experimental period (treatment regimens at each time point align with those described in Figure 1A).

### Gut microbiota test

We collected colon of every rat of different group, and squeezed out about 0.2g feces to be used for detecting gut microbiome. Total genome DNA from each feces sample was extracted using CTAB (Cetyltrimethylammonium Bromide) method. Concentration and purity of the DNA was monitored on 1% agarose gels. According to the concentration, the DNA was diluted to 1ng/μL using sterile water. 16SrRNA gene of distinct regions (16S V4/16S V3/16S V3-V4/16S V4-V5) was amplified used specific primers (table 1) with the barcode. PCR reaction was carried out with 15μL of Phusion® High-Fidelity PCR Master Mix (New England Biolabs), 2μM forward and reverse primers, and 10ng template DNA. Thermal cycling consisted of 98℃ for 1min, followed by 30 cycles of 98℃ for 10s, 50℃ for 30s, and 72℃ for 30s. Finally 72℃ for 5min.

**Table 1.**
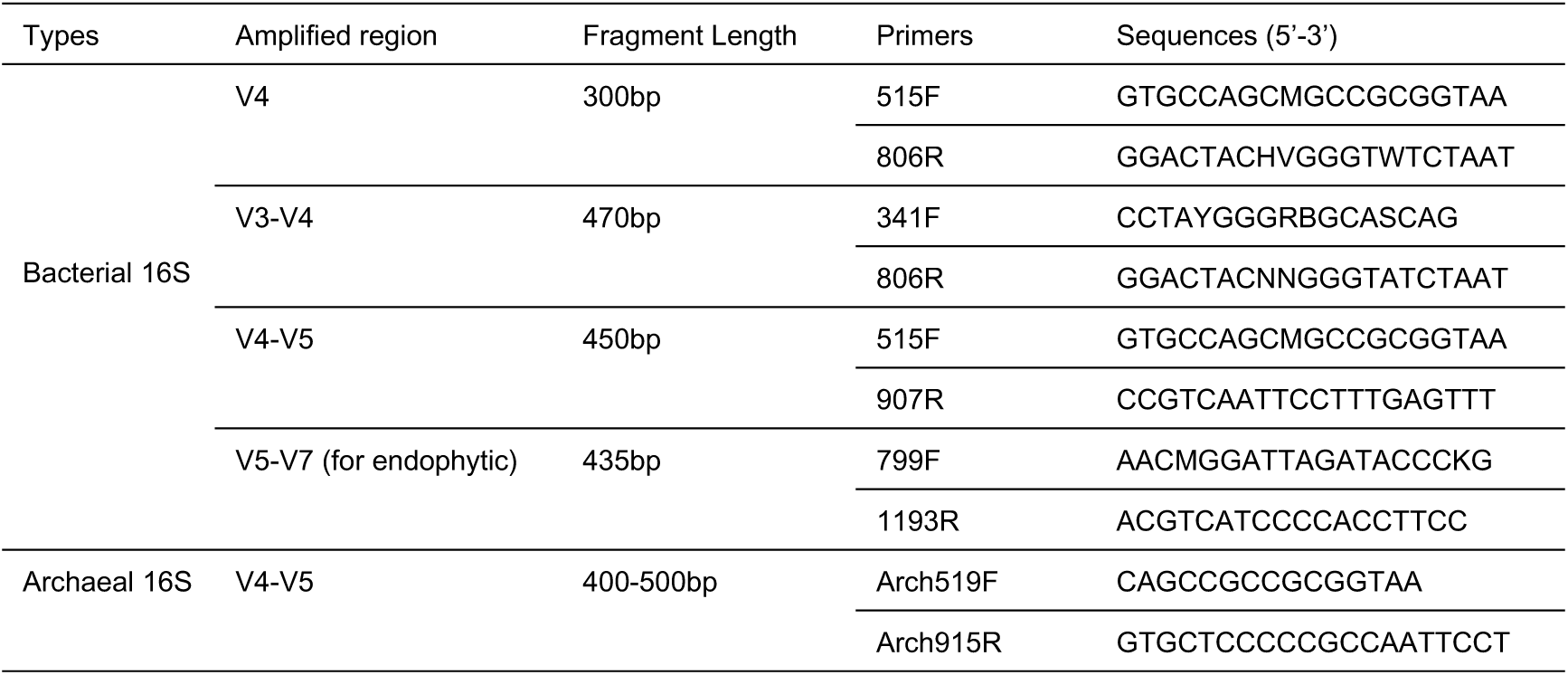
primers used for amplifying 16SrRNA gene of distinct regions.

The PCR products were mixed with same volume of 1X loading buffer (contained SYBR Green) and detected on 2% agarose gel by electrophoresis. Then, the PCR products were purified with Qiagen Gel Extraction Kit (Qiagen, Germany). DNA sequencing libraries were generated using TruSeq® DNA PCR-Free Sample Preparation Kit (Illumina, USA) following manufacturer’s recommendations and index codes were added. The libraries quality was assessed on the Qubit@ 2.0 Fluorometer (Thermo Scientific) and Agilent Bioanalyzer 2100 system. At last, the libraries were sequenced on an Illumina NovaSeq platform and 250 bp paired-end reads were generated. Then the paired-end reads were assigned to samples based on their unique barcodes and truncated by cutting off the barcodes and primer sequences.

According to the fastp (v0.22.0, https://github.com/OpenGene/fastp), the raw tags were filtered under specific filtering conditions to obtain high quality clean tags. Paired-end reads were merged using FLASH (v1.2.11, http://ccb.jhu.edu/software/FLASH/)(14), when at least some of the reads overlap the read generated from the opposite end of the same DNA fragment merge paired-end reads, paired-end reads were merged. Compared the tag with the reference database (Silva database (16S/18S) https://www.arb-silva.de/) using UCHIME Algorithm (http://www.drive5.com/usearch/manual/uchime_algo.html) (15) to discover chimera sequences and removed them(16). Then the Effective Tags were finally obtained.

Amplicon sequence variants (ASV) were analyzed by Deblur, which used error profiles to obtain putative error-free sequences from Illumina sequencing platform. Each representative sequence was annotated taxonomic information, using Silva Database (http://www.arb-silva.de/) (17) based on Mothur algorithm.

In order to study phylogenetic relationship of different ASVs, and the difference of the dominant species in different samples (groups), multiple sequence alignment was conducted using the MAFFT (v7.490, https://mafft.cbrc.jp/alignment/software/). ASVs abundance were normalized using a standard of sequence number corresponding to the sample with the least sequences. Subsequent analysis of alpha diversity and beta diversity were all performed basing on this output normalized data. Alpha diversity was applied in analyzing complexity of species diversity for a sample through Chao1 index, this index was calculated with QIIME and displayed with R software (Version 4.1.2). Beta diversity analysis was used to evaluate differences of samples in species complexity, Beta diversity on both weighted and unweighted unifrac were calculated by QIIME software. Using Non-Metric Multi-Dimensional Scaling (NMDS) to reflected the difference between samples.

Basing on KEGG (Kyoto Encyclopedia of Genes and Genomes) database, Using PICRUSt2 (Phylogenetic Investigation of Communities by Reconstruction of Unobserved States) (18) to forecast functions of 16S rDNA.

### Metabolites test

We used LC/MS (Liquid Chromatograph/Mass Spectrometer) method to untargeted metabolomics test and analyze the metabolites from feces of mice.

#### 1 Sample preparation and extraction

Fecal samples stored at –80 °C refrigerator were thawed on ice. Every 20 mg sample was added 400 μL solution (Methanol: Water = 7:3, V/V) containing internal standard, then vortexed for 3 min. After blending, we sonicated the samples in an ice bath for 10 min and vortexed it for 1 min, then placed them in –20 °C for 30 min. Next, we centrifuged the samples at 12000 rpm for 10 min in 4 °C. And then the sediments were removed, the supernatants were centrifuged at 12000 rpm for 3 min in 4 °C. In the end, a 200 μL aliquots of supernatant were transferred for LC-MS analysis.

#### 2 HPLC (High Performance Liquid Chromatography) Conditions

Two LC/MS methods were employed to test samples. A aliquot was analyzed using positive ion conditions and eluted from T3 column (Waters ACQUITY Premier HSS T3 Column 1.8 µm, 2.1 mm * 100 mm) using 0.1 % formic acid in water (solvent A) and 0.1 % formic acid in acetonitrile (solvent B) in the following gradient: 5 to 20 % in 2 min, 60 % in 3 mins, 99 % in 1 min and held for 1.5 min, then come back to 5 % mobile phase B within 0.1 min, held for 2.4 min. The analytical conditions were as follows: 40 °C (column temperature); 0.4 mL/min (flow rate); 4 μL (injection volume); Another aliquot was using negative ion conditions was the same as the elution gradient of positive mode.

#### 3 MS Conditions (AB)

The Analyst TF 1.7.1 Software (Sciex, Concord, ON, Canada) was employed to operate the data acquisition using the information-dependent acquisition (IDA) mode. The source parameters were set as follows: ion source gas 1 (GAS1), 50 psi; ion source gas 2 (GAS2), 50 psi; curtain gas (CUR), 25 psi; temperature(TEM), 550 °C; declustering potential (DP), 60 V, or−60 V in positive or negative modes, respectively; and ion spray voltage floating (ISVF), 5000 V or−4000 V in positive or negative modes, respectively. The TOF MS scan parameters were set as follows: mass range, 50–1000 Da; accumulation time, 200 ms; and dynamic background subtract, on. The product ion scan parameters were set as follows: mass range, 25-1000 Da; accumulation time, 40 ms; collision energy, 30 or –30 V in positive or negative modes, respectively; collision energy spread, 15; resolution, UNIT; charge state, 1 to 1; intensity, 100 cps; exclude isotopes within 4 Da; mass tolerance, 50 ppm; maximum number of candidate ions to monitor per cycle, 18.

#### 4 Analytical methods

The original data file from LC-MS was converted into format mzXML using ProteoWizard software. The XCMS program was used to perform the peak extraction, peak alignment and retention time correctio. The peak area was corrected using the “SVR” method. The peaks with detetion rate lower than 50 % in each group of samples were discarded. After that, metabolic identification information was obtained by searching the laboratory’s self-built database, integrated public database, AI database and metDNA.

Unsupervised PCA (principal component analysis) was performed by using R (www.r-project.org). The data was unit variance scaled before unsupervised PCA. The HCA (hierarchical cluster analysis) results of samples and metabolites were presented as heatmaps with dendrograms. Both HCA and PCC were carried out by R package Complex Heat map. For HCA, normalized signal intensities of metabolites (unit variance scaling) are visualized as a color spectrum.

#### 5 Differential metabolites selected

For two-group analysis, differential metabolites were determined by VIP (VIP > 1) and *P*-value (*P*-value < 0.05, Student’s t test). VIP values were extracted from OPLS-DA result, which also contain score plots and permutation plots, was generated using R package MetaboAnalystR. The data was log transform (log2) and mean centering before OPLS-DA.

In order to avoid overfitting, a permutation test (200 permutations) was performed. 6 KEGG annotation and enrichment analysis We used the KEGG Compound database (http://www.kegg.jp/kegg/compound/) to annotate the identified metabolites then mapped the annotated metabolites to KEGG Pathway database (http://www.kegg.jp/kegg/pathway.html). Significantly enriched pathways are identified with a hypergeometric test’s *P*-value for a given list of metabolites.

### Statistical software and basic methods

Except R software (Version 4.1.2), excel2019 and SPSS26.0 was also employed to collect and analyze some data. Comparing between more than 2 groups, normal distribution and homoscedasticity data use one-way ANOVA (Analysis of Variance), abnormal distribution or heterogeneity of variance data use Kruskal-Wallis test (K-W test/H test).

## Results

### FPG and weight

Body weights among the 9 groups were comparable at the start of the experiment (*H*=6.846, *P*=0.553). Prior to STZ administration, the mean FPG levels of all groups remained stable around 5 mmol/L. Following intraperitoneal STZ injection, FPG levels increased sharply within 5 days but subsequently declined over the final 3 weeks (Figure 1A). At the end of the experiment, body weights of the RFI, bT3I, MI, and HFI groups were significantly lower than those of the CF group (*F*=6.695, *P*=0.01) (Figure 1B). Notably, except for the HFI group, FPG levels in the other four groups did not differ significantly. Although the RFI, bT3I, and MI treatment groups alleviated hyperglycemia in T2D rats, these interventions failed to halt the progressive weight loss. Detailed data on individual sample FPG levels and body weights are provided in Supplementary Table 1.

### Gut microbial diversity

We characterized the diversity of ASVs across the 9 experimental groups to reflect intergroup differences in the gut microbiome. Our results revealed that the BL group exhibited the lowest ASV richness, whereas the CF displayed the highest ASV count. When rats developed T2D, their ASV abundance decreased compared to age-matched control rats (CT group). Continuous feeding with TMF further reduced ASV richness in T2D rats. Notably, switching to MF or combining MF with MET reversed this decline. Collectively, these findings suggest that unhealthy dietary patterns reduce gut microbiota diversity, but which can be restored by adopting a healthy diet or incorporating antidiabetic therapeutics (e.g., MET) instead of continuing an unhealthy diet.

Comparisons of both shared and unique ASVs across groups indicated that younger rats (BL group) harbored a greater number of unique ASVs (Figure 2A). The CF group consistently exhibited the highest α-diversity, underscoring that a healthy diet promotes high gut microbial diversity. In contrast, the BL, T2D, HFI, and Death groups displayed the lowest α-diversity. These observations led us to conclude that reduced gut microbial diversity is associated with three key stages: (1) early life (young age); (2) the T2D pathological state; and (3) the terminal stage (death). When T2D rats were transitioned to MF (RFI group), MF + β-T3 (bT3I group), or MF + MET (MI group) instead of TMF, their α-diversity significantly increased. Notably, RFI and bT3I groups recovered to α-diversity levels comparable to CF, whereas the MI group remained slightly lower than CF. Overall, sustained feeding with MF promoted gradual increases in gut microbial diversity in adult rats (BL< CT < CF). Conversely, a high-fat diet (TMF) maintained low diversity until death, with no significant differences observed between BL, T2D, HFI, and Death groups. Intriguingly, switching from a high-fat to a standard diet restored diversity to optimal levels, even when supplemented with low-dose β-T3. However, MET appeared to impede the recovery of microbial diversity when rats were transitioned to MF (Figure 2B). Additionally, while β-T3 did not increase ASV richness, it did not hinder diversity recovery, suggesting that β-T3 may facilitate the proliferation of specific homologous bacterial taxa through its stimulatory effects.

**Figure 2.**
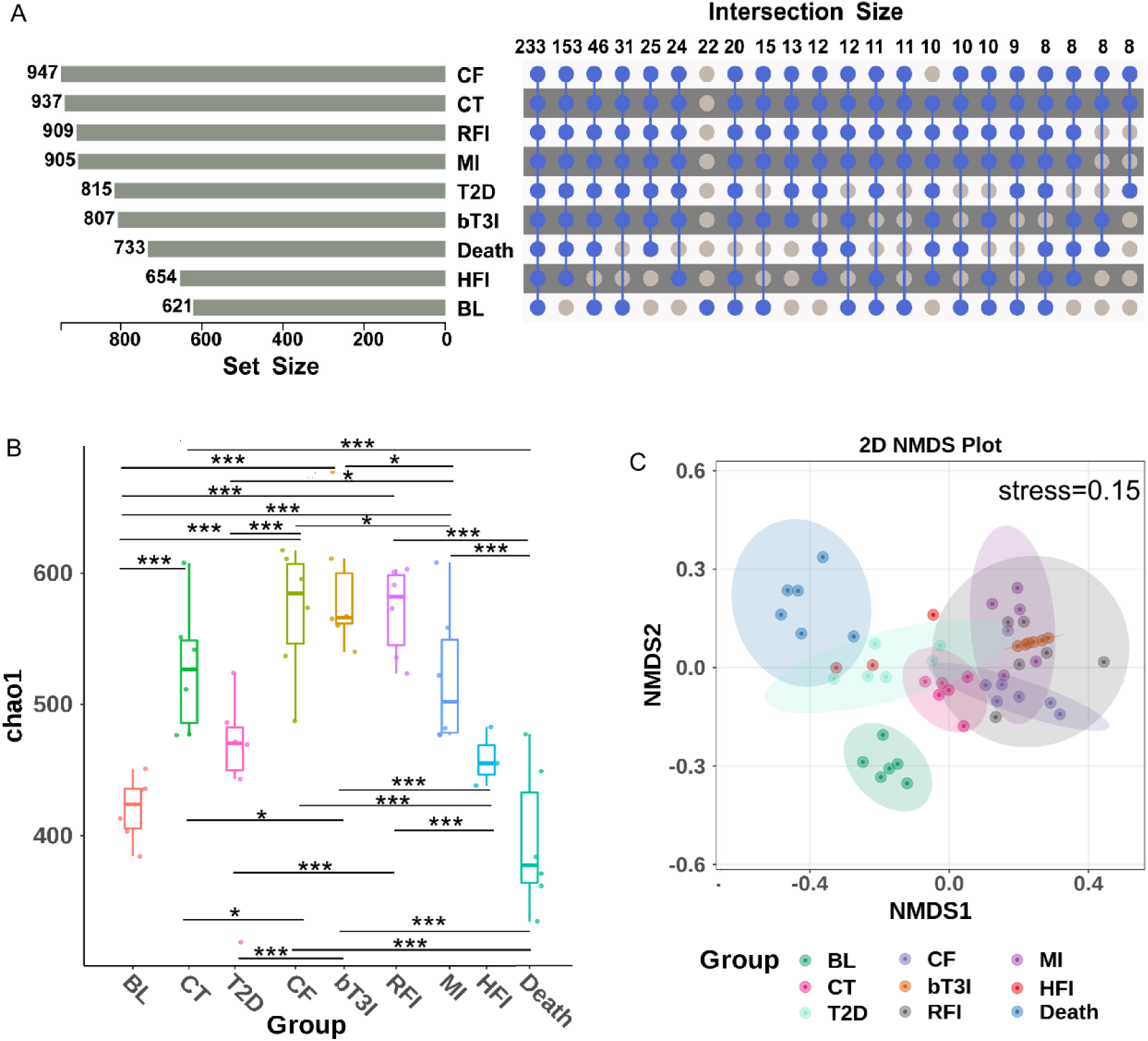
Gut microbial diversity across the 9 groups. A. Venn diagram of ASVs among the 9 groups. The left panel shows the total ASV count per group, while the right panel illustrates shared and unique ASVs between groups. Blue markers denote ASVs present in the group (corresponding to the left panel), and gray markers indicate ASVs absent from the group. B. α-diversity analysis (Chao1 index) using the Kruskal-Wallis test. *Indicates significant differences between groups connected by a horizontal line.C. β-diversity visualization (Bray-Curtis dissimilarity) via non-metric multidimensional scaling (NMDS) ordination. The stress value is 0.15 (<0.2), indicating the results reliably capture intergroup differences.

Intergroup β-diversity analysis revealed significant differences, particularly between BL, T2D, and Death groups, which clustered far from other groups. This pattern indicates that these three stages harbor distinct gut microbiota signatures (Figure 2C).

### Succession of gut microbiota

Differential relative abundances across six microbial taxonomic levels (phylum, class, order, family, genus, and species) were analyzed to characterize the successional patterns of gut microbiota during T2D progression (Figure 3). At the phylum level, p Firmicutes and p Bacteroidota emerged as the dominant taxa. In healthy young rats, Firmicutes exhibited the highest relative abundance, which gradually decreased with age (BL > CT > CF). Notably, the relative abundance of p Campylobacterota increased significantly in the CF group, potentially linked to age-related physiological changes or hormonal regulation. In T2D rats, however, p Firmicutes abundance was markedly elevated compared to control groups (CT and CF), and this trend persisted with continuous TMF feeding (HFI group). By the terminal stage (Death), both p Firmicutes and p Bacteroidota abundances declined, whereas p Proteobacteria and p Actinobacteria increased conspicuously, p Proteobacteria even emerged as a dominant taxon, reaching abundances comparable to p Bacteroidota.

**Figure 3.**
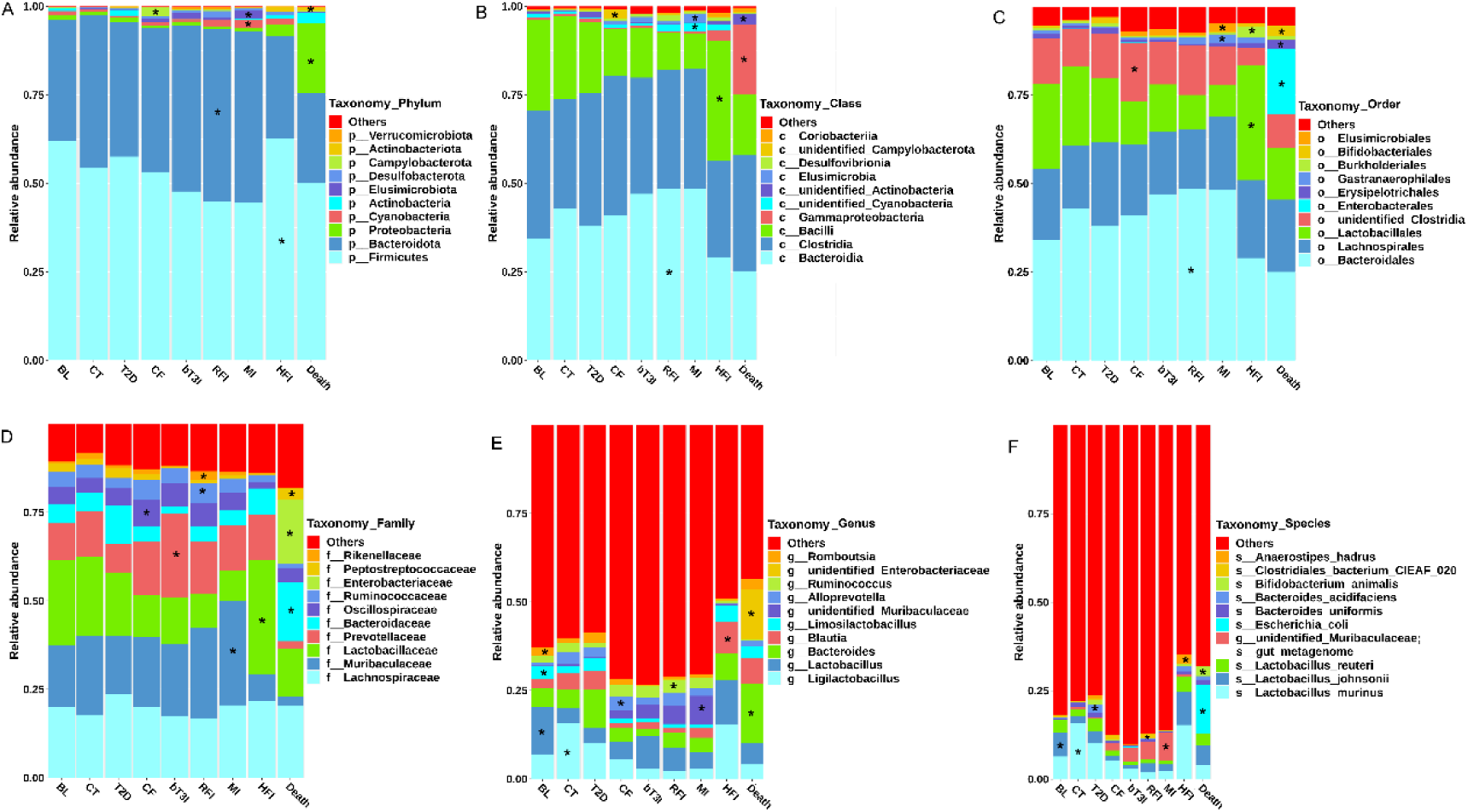
Percent stacked bar charts characterize the successional dynamics of gut microorganisms across six taxonomic levels (phylum, class, order, family, genus, and species). The x-axis denotes distinct experimental groups, while the y-axis represents the relative abundance of microbial taxa at each level. An asterisk (*) indicates microbial taxa identified as biomarkers specific to a given group. Subplots are structured as follows: A. Phylum-level microbial changes; B. Class-level microbial changes; C. Order-level microbial changes; D. Family-level microbial changes; E. Genus-level microbial changes; F. Species-level microbial changes.

Intriguingly, when T2D rats were transitioned to MF (RFI group), or received MF supplemented with β-T3 (bT3I group) or MET (MI group) instead of TMF, the relative abundances of these two dominant taxa (Firmicutes and Bacteroidota) recovered to levels approximating the control group (CF). Notably, β-T3 supplementation (bT3I) more effectively restored microbial balance, whereas MF alone primarily increased p Bacteroidota abundance. Conversely, MET treatment elevated p Cyanobacteria and p Elusimicrobiota.

In the rest of other microbial levels, class, order, family, genus and species, the dominant bacteria showed specific change rule, respectively. And all the 6 levels, RFI, bT3I and MI three groups exhibited the same trend with that of CF. That means all the three measures could help the disease mice restore their gut bacteria. Using LEfSe (LDA≥4, P> 0.05)(19), we identified the statistically different biomarkers in each group (Table 2). In the species levels. the *g Lactobacillus, g Limosilactobacillus* and *g Romboutsia* three genera of Firmicutes play key roles in the early age (BL), and the *s Lactobacillus_johnsonii* of *g Lactobacillus* may be the key species. When rats are 4 months old, *g_Ligllactobacillus* and *s_ Ligllactobacillus murinus* of *g Ligilactobacillus* become the key factors instead of *g Lactobacillus*. Following the rising of age, the gut microbiota became more balanced, the number of dominant microbial species is fewer than that of early age, and the disparities between the differently dominant microorgnisms reduced. *g Alloprevotella* in CF is higher than other groups in abundance, that means the genus would expand bit by bit in the life progress with eating healthy food. And when we had fed the rats with TMF for 8 weeks and induced them to T2D models, we found the *B. acidifaciens* had a transitory dilation, that probably because of the compensatory mechanism. The T2D rats were fed by TMF for four weeks, the number of the *g Blautia*, and *s_Anaerostipes_hadrus* (*A. hadrus*) increased and they became the biomarkers of HFI. Until the T2D rats dead because of eating fat food for couple of weeks, *E. coli* becomes the dominant microorganism, the *E. coli* and *s_Bifidobacterium_animalis* (*B. animalis*) could be the biomarkers of death mice in the species level. In the three intervention groups, RFI gave rise to population increase of *g Ruminococcus* and *s_Clostridiales_bacterium*_CIEAF_020, MI made *g unidentified_Muribaculaceae* and *s_gut_metagenome* of this genus higher in number, that means the two genera probably are the main bacteria to metabolize MET. What impressed us was the β-T3 & MF feeding rats showed the more similar to the gut microbiota of CF group in genus and species levels. That can be explained again the β-T3 is a beneficial factor to regulate T2D.

**Table 2.**
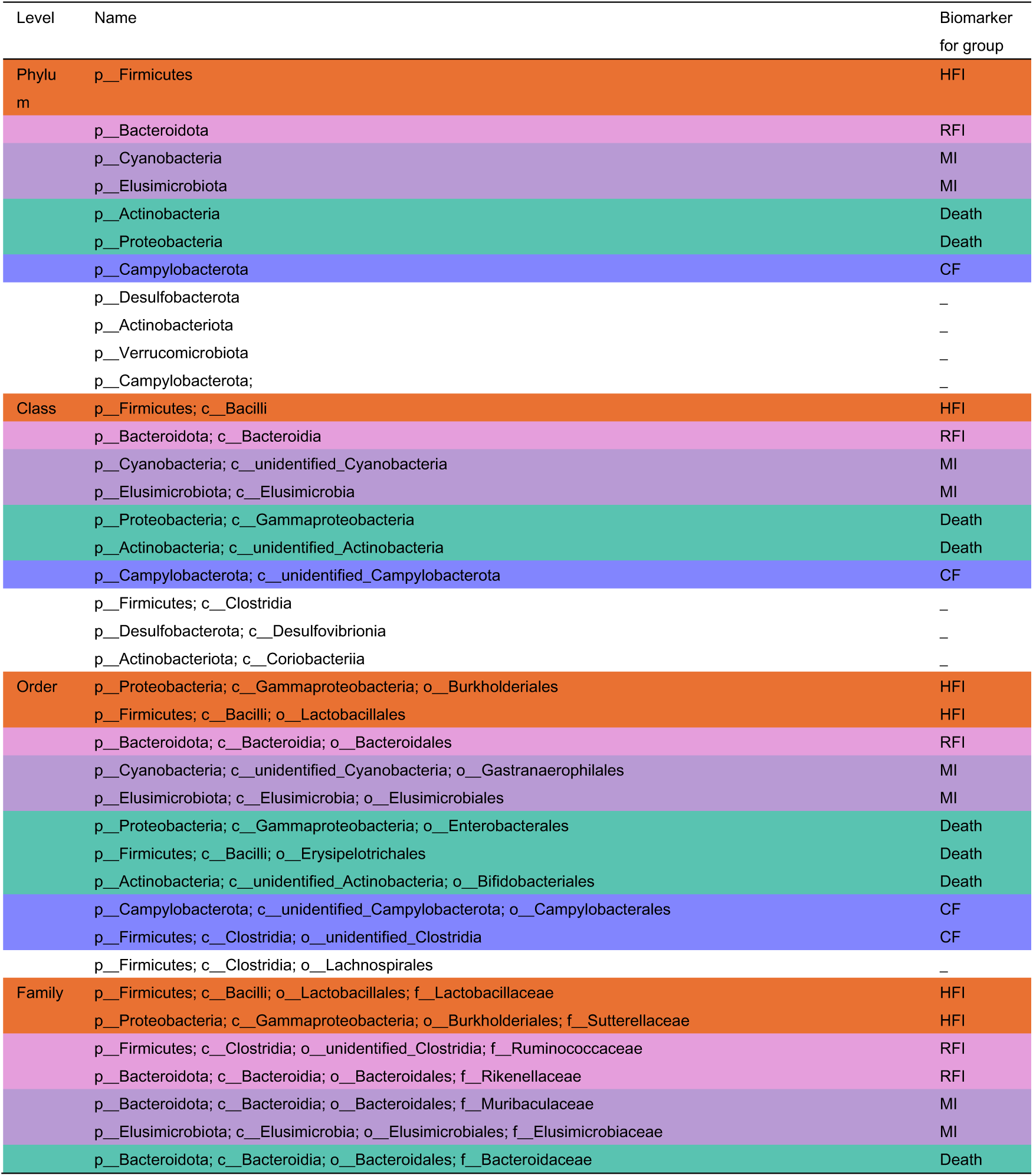
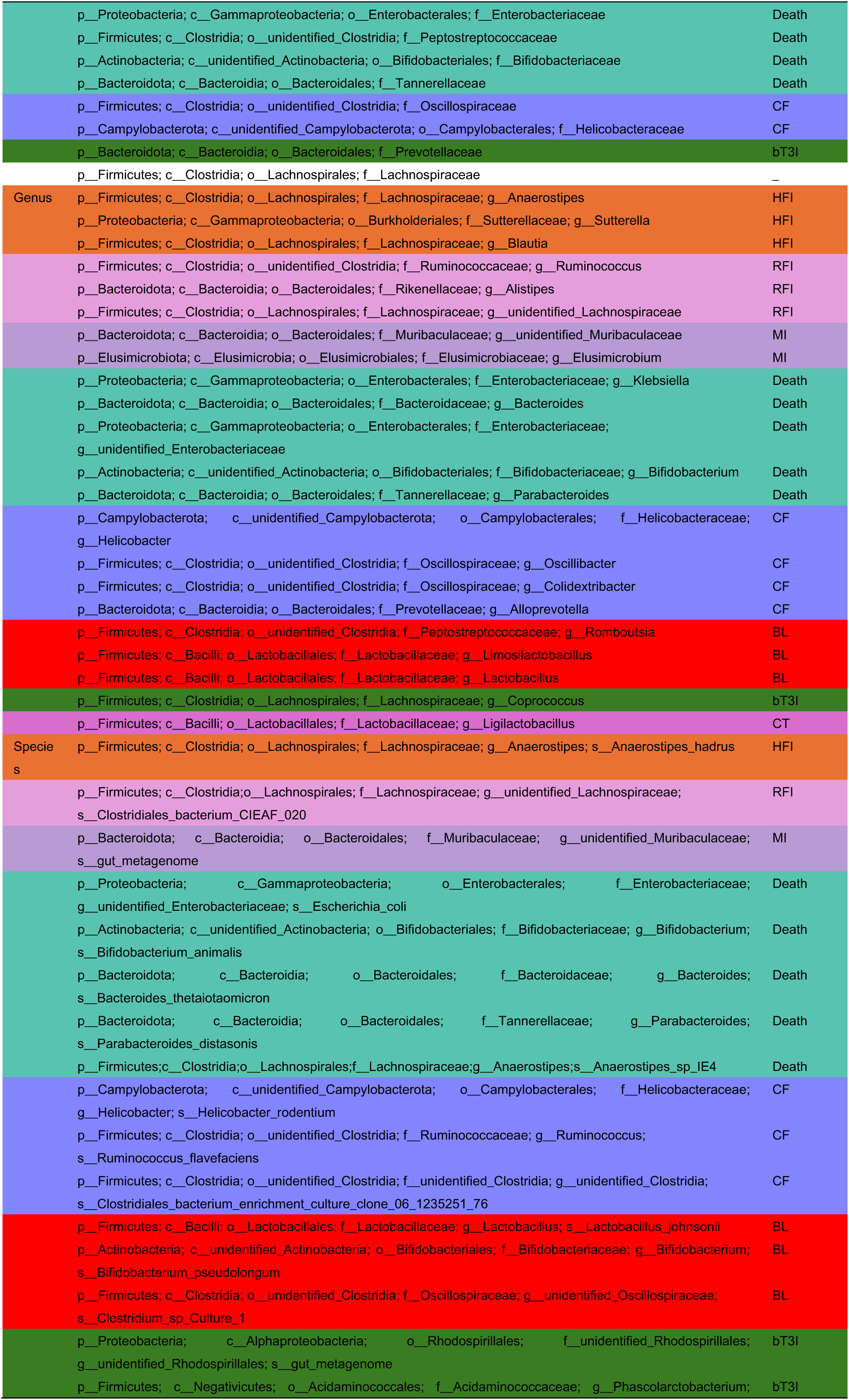
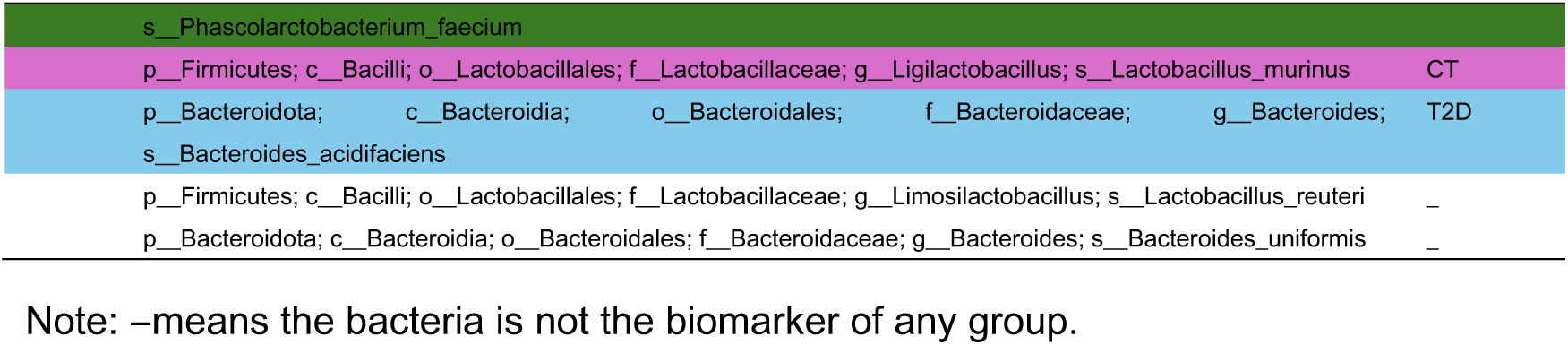
Phylogenetic relationships of dominant microbial species and biomarker information across different groups.

| Level | Name | Biomarker<br>for group |
| --- | --- | --- |
| Phylum | p__Firmicutes | HFI |
|  | p__Bacteroidota | RFI |
|  | p__Cyanobacteria | MI |
|  | p__Elusimicrobiota | MI |
|  | p__Actinobacteria | Death |
|  | p__Proteobacteria | Death |
|  | p__Campylobacterota | CF |
|  | p__Desulfobacterota | – |
|  | p__Actinobacteriota | – |
|  | p__Verrucomicrobiota | – |
|  | p__Campylobacterota; | – |
| Class | p__Firmicutes; c__Bacilli | HFI |
|  | p__Bacteroidota; c__Bacteroidia | RFI |
|  | p__Cyanobacteria; c__unidentified_Cyanobacteria | MI |
|  | p__Elusimicrobiota; c__Elusimicrobia | MI |
|  | p__Proteobacteria; c__Gammaproteobacteria | Death |
|  | p__Actinobacteria; c__unidentified_Actinobacteria | Death |
|  | p__Campylobacterota; c__unidentified_Campylobacterota | CF |
|  | p__Firmicutes; c__Clostridia | – |
|  | p__Desulfobacterota; c__Desulfovibrionia | – |
|  | p__Actinobacteriota; c__Coriobacteriia | – |
| Order | p__Proteobacteria; c__Gammaproteobacteria; o__Burkholderiales | HFI |
|  | p__Firmicutes; c__Bacilli; o__Lactobacillales | HFI |
|  | p__Bacteroidota; c__Bacteroidia; o__Bacteroidales | RFI |
|  | p__Cyanobacteria; c__unidentified_Cyanobacteria; o__Gastranaerophilales | MI |
|  | p__Elusimicrobiota; c__Elusimicrobia; o__Elusimicrobiales | MI |
|  | p__Proteobacteria; c__Gammaproteobacteria; o__Enterobacterales | Death |
|  | p__Firmicutes; c__Bacilli; o__Erysipelotrichales | Death |
|  | p__Actinobacteria; c__unidentified_Actinobacteria; o__Bifidobacteriales | Death |
|  | p__Campylobacterota; c__unidentified_Campylobacterota; o__Campylobacterales | CF |
|  | p__Firmicutes; c__Clostridia; o__unidentified_Clostridia | CF |
|  | p__Firmicutes; c__Clostridia; o__Lachnospirales | – |
| Family | p__Firmicutes; c__Bacilli; o__Lactobacillales; f__Lactobacillaceae | HFI |
|  | p__Proteobacteria; c__Gammaproteobacteria; o__Burkholderiales; f__Sutterellaceae | HFI |
|  | p__Firmicutes; c__Clostridia; o__unidentified_Clostridia; f__Ruminococcaceae | RFI |
|  | p__Bacteroidota; c__Bacteroidia; o__Bacteroidales; f__Rikenellaceae | RFI |
|  | p__Bacteroidota; c__Bacteroidia; o__Bacteroidales; f__Muribaculaceae | MI |
|  | p__Elusimicrobiota; c__Elusimicrobia; o__Elusimicrobiales; f__Elusimicrobiaceae | MI |
|  | p__Bacteroidota; c__Bacteroidia; o__Bacteroidales; f__Bacteroidaceae | Death |
|  | p__Proteobacteria; c__Gammaproteobacteria; o__Enterobacterales; f__Enterobacteriaceae | Death |
|  | p__Firmicutes; c__Clostridia; o__unidentified_Clostridia; f__Peptostreptococcaceae | Death |
|  | p__Actinobacteria; c__unidentified_Actinobacteria; o__Bifidobacteriales; f__Bifidobacteriaceae | Death |
|  | p__Bacteroidota; c__Bacteroidia; o__Bacteroidales; f__Tannerellaceae | Death |
|  | p__Firmicutes; c__Clostridia; o__unidentified_Clostridia; f__Oscillospiraceae | CF |
|  | p__Campylobacterota; c__unidentified_Campylobacterota; o__Campylobacterales; f__Helicobacteraceae | CF |
|  | p__Bacteroidota; c__Bacteroidia; o__Bacteroidales; f__Prevotellaceae | bT3I |
|  | p__Firmicutes; c__Clostridia; o__Lachnospirales; f__Lachnospiraceae | - |
| Genus | p__Firmicutes; c__Clostridia; o__Lachnospirales; f__Lachnospiraceae; g__Anaerostipes | HFI |
|  | p__Proteobacteria; c__Gammaproteobacteria; o__Burkholderiales; f__Sutterellaceae; g__Sutterella | HFI |
|  | p__Firmicutes; c__Clostridia; o__Lachnospirales; f__Lachnospiraceae; g__Blautia | HFI |
|  | p__Firmicutes; c__Clostridia; o__unidentified_Clostridia; f__Ruminococcaceae; g__Ruminococcus | RFI |
|  | p__Bacteroidota; c__Bacteroidia; o__Bacteroidales; f__Rikenellaceae; g__Alistipes | RFI |
|  | p__Firmicutes; c__Clostridia; o__Lachnospirales; f__Lachnospiraceae; g__unidentified_Lachnospiraceae | RFI |
|  | p__Bacteroidota; c__Bacteroidia; o__Bacteroidales; f__Muribaculaceae; g__unidentified_Muribaculaceae | MI |
|  | p__Elusimicrobiota; c__Elusimicrobia; o__Elusimicrobiales; f__Elusimicrobiaceae; g__Elusimicrobium | MI |
|  | p__Proteobacteria; c__Gammaproteobacteria; o__Enterobacterales; f__Enterobacteriaceae; g__Klebsiella | Death |
|  | p__Bacteroidota; c__Bacteroidia; o__Bacteroidales; f__Bacteroidaceae; g__Bacteroides | Death |
|  | p__Proteobacteria; c__Gammaproteobacteria; o__Enterobacterales; f__Enterobacteriaceae; g__unidentified_Enterobacteriaceae | Death |
|  | p__Actinobacteria; c__unidentified_Actinobacteria; o__Bifidobacteriales; f__Bifidobacteriaceae; g__Bifidobacterium | Death |
|  | p__Bacteroidota; c__Bacteroidia; o__Bacteroidales; f__Tannerellaceae; g__Parabacteroides | Death |
|  | p__Campylobacterota; c__unidentified_Campylobacterota; o__Campylobacterales; f__Helicobacteraceae; g__Helicobacter | CF |
|  | p__Firmicutes; c__Clostridia; o__unidentified_Clostridia; f__Oscillospiraceae; g__Oscillibacter | CF |
|  | p__Firmicutes; c__Clostridia; o__unidentified_Clostridia; f__Oscillospiraceae; g__Colidextribacter | CF |
|  | p__Bacteroidota; c__Bacteroidia; o__Bacteroidales; f__Prevotellaceae; g__Alloprevotella | CF |
|  | p__Firmicutes; c__Clostridia; o__unidentified_Clostridia; f__Peptostreptococcaceae; g__Romboutsia | BL |
|  | p__Firmicutes; c__Bacilli; o__Lactobacillales; f__Lactobacillaceae; g__Limosilactobacillus | BL |
|  | p__Firmicutes; c__Bacilli; o__Lactobacillales; f__Lactobacillaceae; g__Lactobacillus | BL |
|  | p__Firmicutes; c__Clostridia; o__Lachnospirales; f__Lachnospiraceae; g__Coprococcus | bT3I |
|  | p__Firmicutes; c__Bacilli; o__Lactobacillales; f__Lactobacillaceae; g__Ligilactobacillus | CT |
| Species | p__Firmicutes; c__Clostridia; o__Lachnospirales; f__Lachnospiraceae; g__Anaerostipes; s__Anaerostipes_hadrus | HFI |
|  | p__Firmicutes; c__Clostridia; o__Lachnospirales; f__Lachnospiraceae; g__unidentified_Lachnospiraceae; s__Clostridiales_bacterium_CIEAF_020 | RFI |
|  | p__Bacteroidota; c__Bacteroidia; o__Bacteroidales; f__Muribaculaceae; g__unidentified_Muribaculaceae; s__gut_metagenome | MI |
|  | p__Proteobacteria; c__Gammaproteobacteria; o__Enterobacterales; f__Enterobacteriaceae; g__unidentified_Enterobacteriaceae; s__Escherichia_coli | Death |
|  | p__Actinobacteria; c__unidentified_Actinobacteria; o__Bifidobacteriales; f__Bifidobacteriaceae; g__Bifidobacterium; s__Bifidobacterium_animalis | Death |
|  | p__Bacteroidota; c__Bacteroidia; o__Bacteroidales; f__Bacteroidaceae; g__Bacteroides; s__Bacteroides_thetaiotaomicron | Death |
|  | p__Bacteroidota; c__Bacteroidia; o__Bacteroidales; f__Tannerellaceae; g__Parabacteroides; s__Parabacteroides_distasonis | Death |
|  | p__Firmicutes; c__Clostridia; o__Lachnospirales; f__Lachnospiraceae; g__Anaerostipes; s__Anaerostipes_sp_IE4 | Death |
|  | p__Campylobacterota; c__unidentified_Campylobacterota; o__Campylobacterales; f__Helicobacteraceae; g__Helicobacter; s__Helicobacter_rodentium | CF |
|  | p__Firmicutes; c__Clostridia; o__unidentified_Clostridia; f__Ruminococcaceae; g__Ruminococcus; s__Ruminococcus_flavefaciens | CF |
|  | p__Firmicutes; c__Clostridia; o__unidentified_Clostridia; f__unidentified_Clostridia; g__unidentified_Clostridia; s__Clostridiales_bacterium_enrichment_culture_clone_06_1235251_76 | CF |
|  | p__Firmicutes; c__Bacilli; o__Lactobacillales; f__Lactobacillaceae; g__Lactobacillus; s__Lactobacillus_johnsonii | BL |
|  | p__Actinobacteria; c__unidentified_Actinobacteria; o__Bifidobacteriales; f__Bifidobacteriaceae; g__Bifidobacterium; s__Bifidobacterium_pseudolongum | BL |
|  | p__Firmicutes; c__Clostridia; o__unidentified_Clostridia; f__Oscillospiraceae; g__unidentified_Oscillospiraceae; s__Clostridium_sp_Culture_1 | BL |
|  | p__Proteobacteria; c__Alphaproteobacteria; o__Rhodospirillales; f__unidentified_Rhodospirillales; g__unidentified_Rhodospirillales; s__gut_metagenome | bT3I |
|  | p__Firmicutes; c__Negativicutes; o__Acidaminococcales; f__Acidaminococcaceae; g__Phascolarctobacterium | bT3I |

|  |  |
| --- | --- |
| s__Phascolarctobacterium_faecium |  |
| p__Firmicutes; c__Bacilli; o__Lactobacillales; f__Lactobacillaceae; g__Ligilactobacillus; s__Lactobacillus_murinus | CT |
| p__Bacteroidota; c__Bacteroidia; o__Bacteroidales; f__Bacteroidaceae; g__Bacteroides; s__Bacteroides_acidifaciens | T2D |
| p__Firmicutes; c__Bacilli; o__Lactobacillales; f__Lactobacillaceae; g__Limosilactobacillus; s__Lactobacillus_reuteri | – |
| p__Bacteroidota; c__Bacteroidia; o__Bacteroidales; f__Bacteroidaceae; g__Bacteroides; s__Bacteroides_uniformis | – |

Note: –means the bacteria is not the biomarker of any group.

### Functional annotation of gut microbial genes

Basing the KEGG database, the functional genes were identified and three levels were all analyzed. We can observe the functional difference between the 9 groups in different levels by Principal coordinate analysis (PCoA), and the death significantly differed from other groups. Meanwhile, there were some specific functional ways existing significant intergroup difference in level 1, level 2 and level 3, respectively (figure 4). There were 6 functions in level 1 having significant intergroup difference, including Genetic Information Processing, Not Included in Pathway or Brite, Environmental Information Processing, Cellular Processes, Human Diseases and Organismal Systems. Although the Metabolism of level 1 did not have statistic otherness between the 9 groups, there were many specific metabolic pathways in level 2 and 3 had obvious difference. In level 2, 49 functions had statistic difference, including genetic information processing, signaling and cellular processes, translation, diseases and so on, 12 of that were metabolism or biosynthesis. We had tested 371 level 3 functional genes, 350 of them had statistic difference between the 9 groups, and about 26.00% in the 350 genes were function genes of metabolism or biosynthesis. The detail information of every samp in three levels was described in supplemental table 2-4.

**Figure 4.**
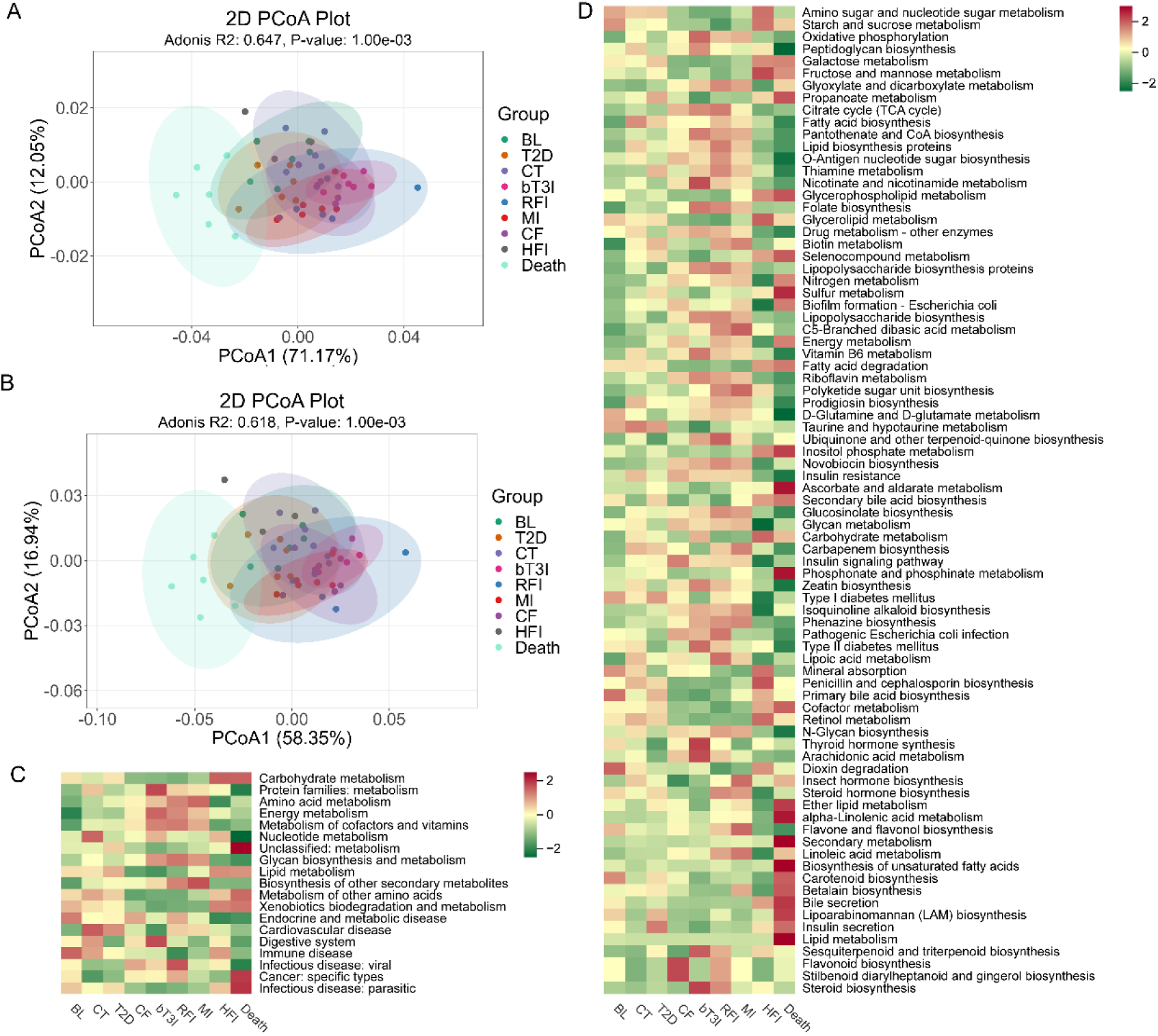
Functional gene comparisons across the 9 groups at different taxonomic levels. A. PCoA ordination results for level 2 functional genes; B. PCoA ordination results for level 3 functional genes; C. All pathways exhibiting significant differences in functional gene abundance between the 9 groups at level 2; D. Metabolic pathways showing significant statistical differences in functional gene abundance between groups at level 3.

Based on the KEGG database, functional genes were annotated, and their abundances were analyzed across three hierarchical levels (Level 1, 2, and 3). PCoA revealed significant functional divergence among the 9 groups, with the Death group exhibiting the most distinct functional profile compared to others (Figure 4). At each level, specific functional pathways displayed marked intergroup differences: Level 1: Six functional categories showed significant intergroup variations, including Genetic Information Processing, Not Included in Pathway or Brite, Environmental Information Processing, Cellular Processes, Human Diseases, and Organismal Systems. Notably, while the Metabolism category at Level 1 did not show overall statistical differences across groups, numerous specific metabolic pathways at Levels 2 and 3 exhibited pronounced variations. Level 2: Forty-nine functions displayed statistical differences, encompassing genetic information processing, signaling, cellular processes, translation, and disease-related pathways. Among these, 12 were associated with metabolism or biosynthesis. Level 3: We analyzed 371 functional genes, of which 350 (94.3%) exhibited statistical differences in abundance across the 9 groups. Approximately 26.0% of these 350 genes were linked to metabolic or biosynthetic functions.

Detailed functional annotations for each sample across the three levels are provided in Supplementary Tables 2–4.

## Metabolites features

A total of 12,920 fecal metabolites were identified across the 9 groups, with 10,129 of these exhibiting significant intergroup differences. Based on the comparison of these 10,129 differential metabolites, a heatmap visually revealed the cluster patterns characteristic of each group (Figure 5). The BL group displayed a distinct profile, with most metabolites downregulated. CT and CF exhibited similar profiles, where most metabolites were balanced in abundance, though a few were significantly upregulated. T2D and HFI shared a comparable pattern, with approximately half of the metabolites upregulated and half downregulated, and only a small fraction remaining balanced. Notably, the three intervention groups bT3I, RFI, and MI clustered closely with CF.

**Figure 5.**
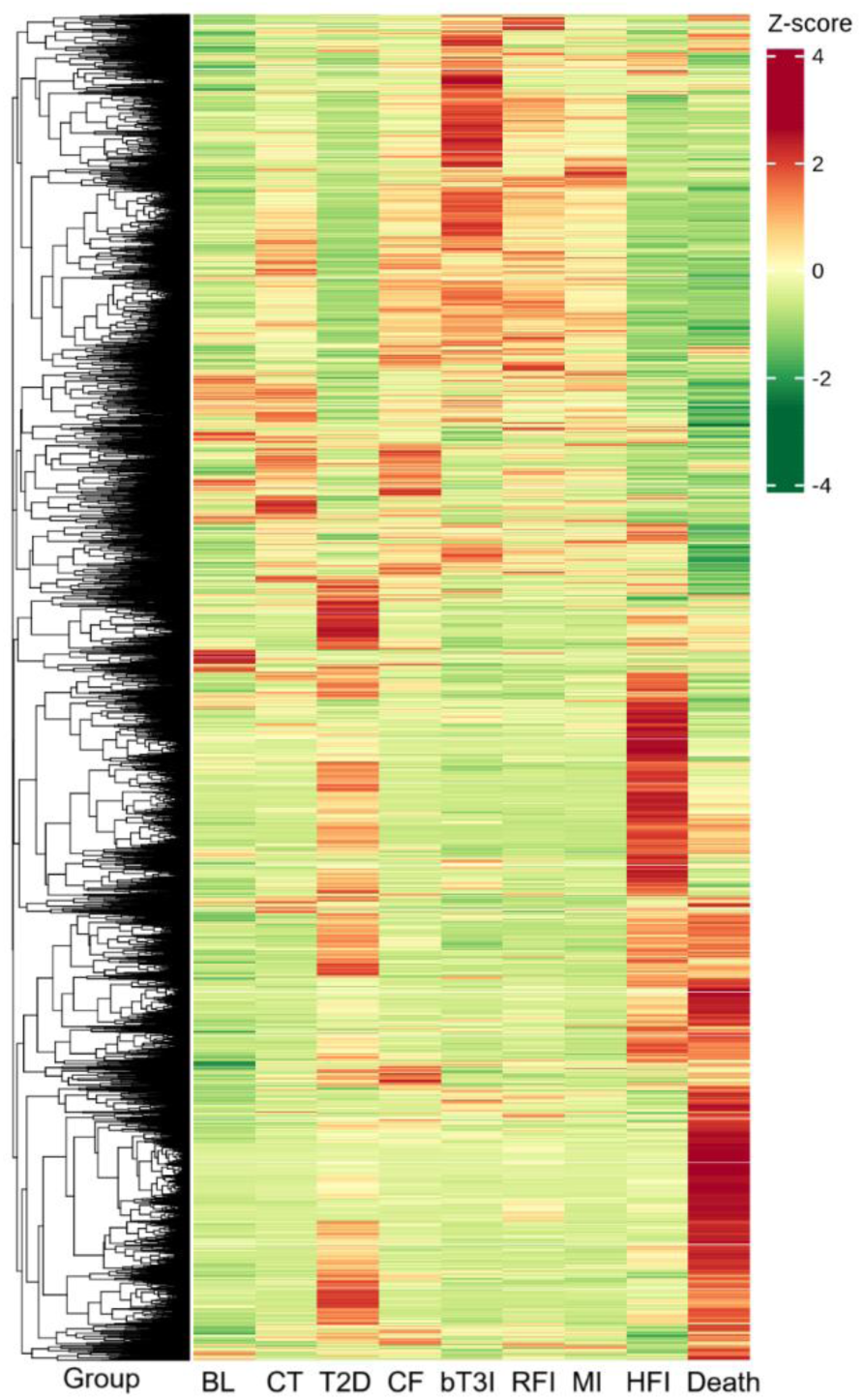
Significantly different metabolites among the 9 groups are color-coded, with red representing upregulated metabolites and green denoting downregulated metabolites.

Of particular interest, certain metabolites were dramatically upregulated in the bT3I group, including benzene and its substituted derivatives, organic acids and their derivatives, heterocyclic compounds, and other metabolites (as confirmed by subsequent analysis). In the Death group, nearly half of the metabolites were dramatically upregulated, and half were downregulated, yet their overall profile resembled that of the T2D group.

We compared two groups and screened statistically significant differentially expressed metabolites from the top 10, identifying BAs, amino acids and their metabolites, as well as organic acids and their derivatives, which exhibited significant upregulation or downregulation in specific groups (Figure 6). Furthermore, comparative analysis of these metabolites across the 9 groups, combined with functional analysis of Level 2 and Level 3 metabolic pathways, confirmed that the metabolism of certain BAs and amino acids serves as a key regulatory pathway in modulating T2D progression.

**Figure 6.**
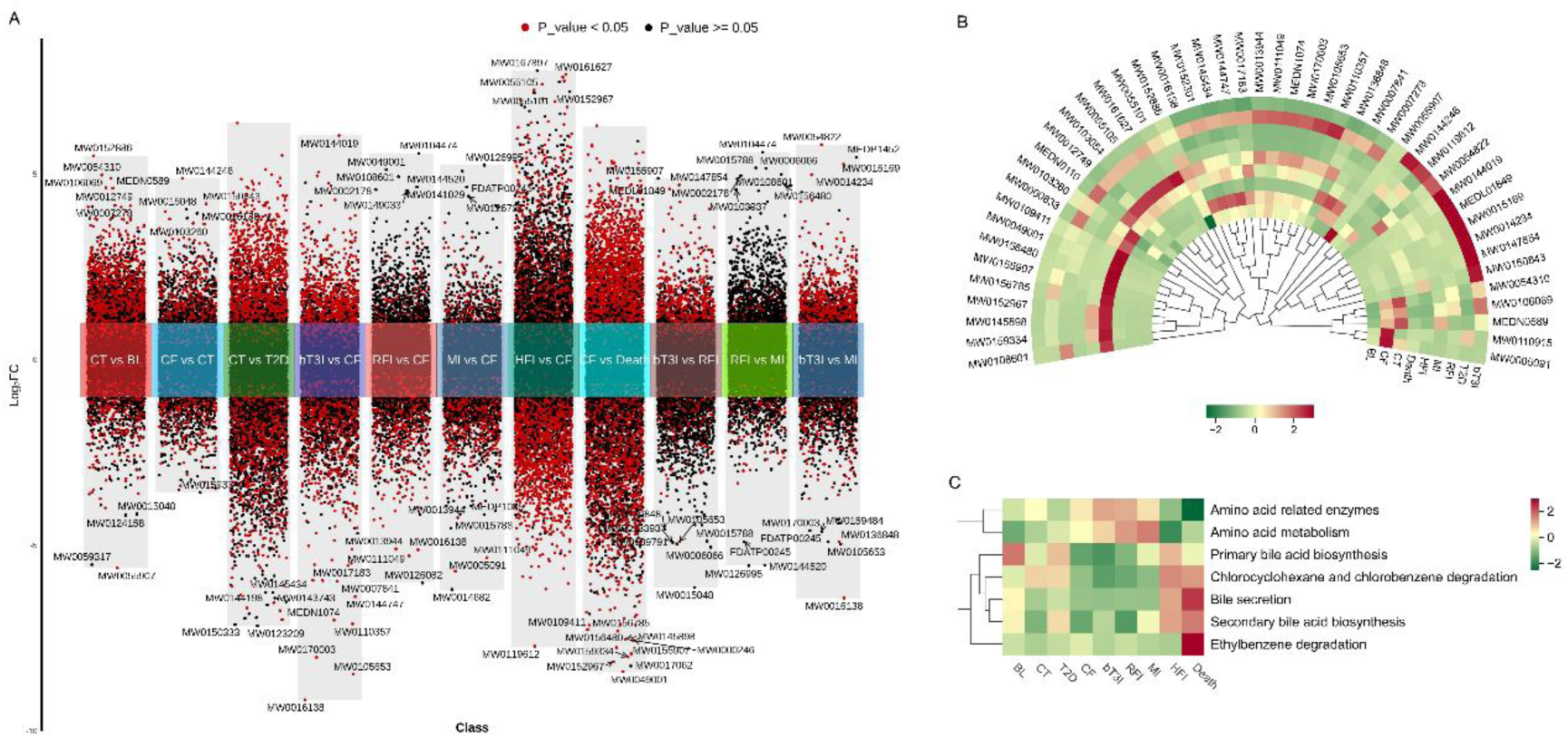
The difference between two groups of gut metabolites. A, Multiple volcano maps (20) of 11 different grouping, the dots that log_2_FC value more or less than 0 means the quantity of the metabolites were up or down in the front group comparing the back group, and the red dots mean the metabolites with statistic difference, the black ones mean no statistically different metabolites. The top 10 log_2_FC value dots were labeled the index of the compounds. B, Heat map of the statistically different compounds from the top 10 comparing between 9 groups. C, Heatmap of comparing metabolic pathways in level 2 and 3 by using gut microbial genes predicting functions.

**Table 3.**
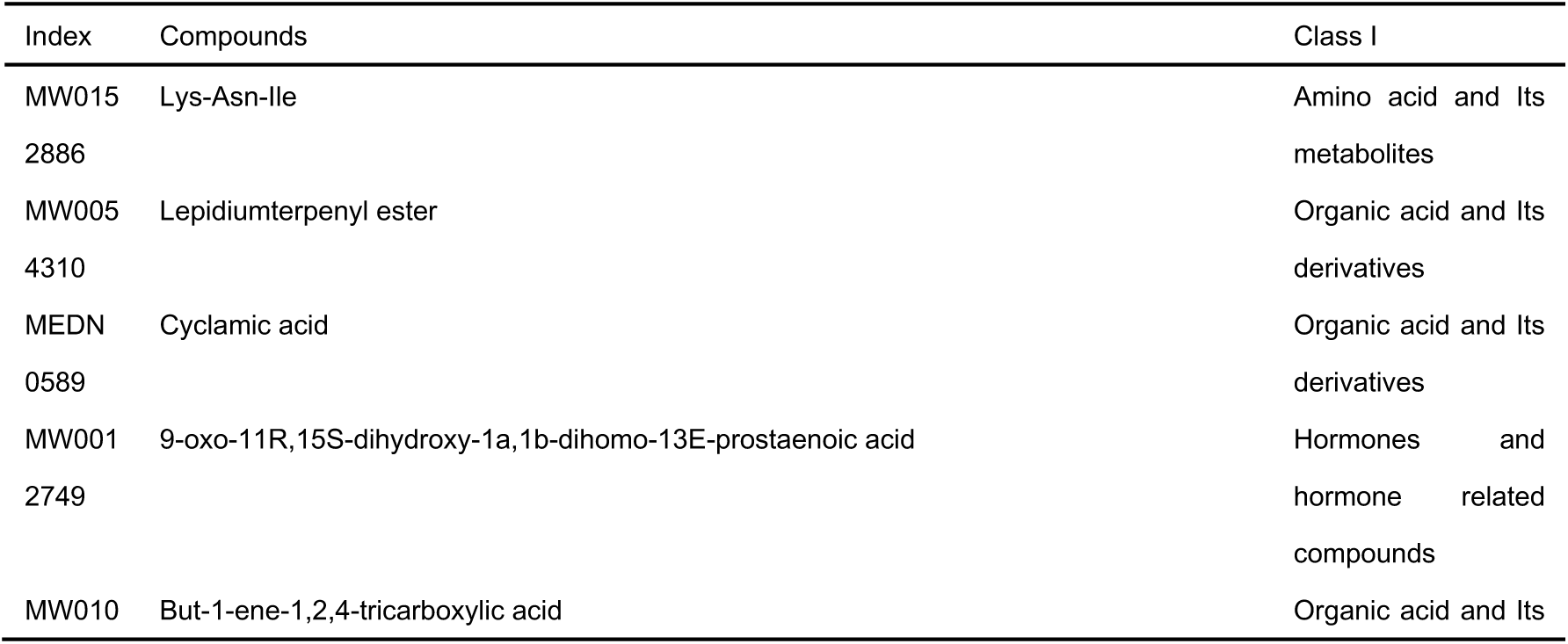
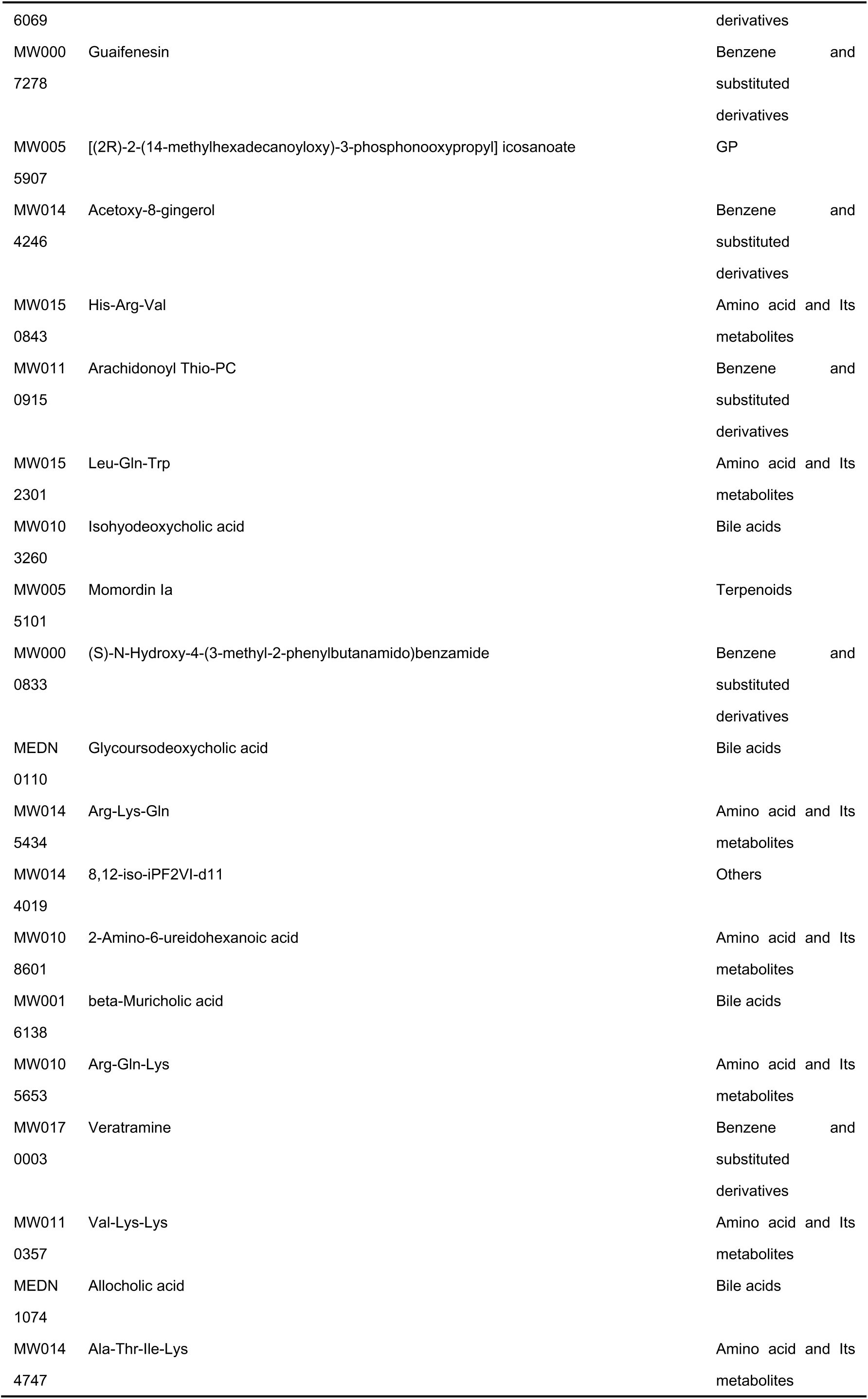
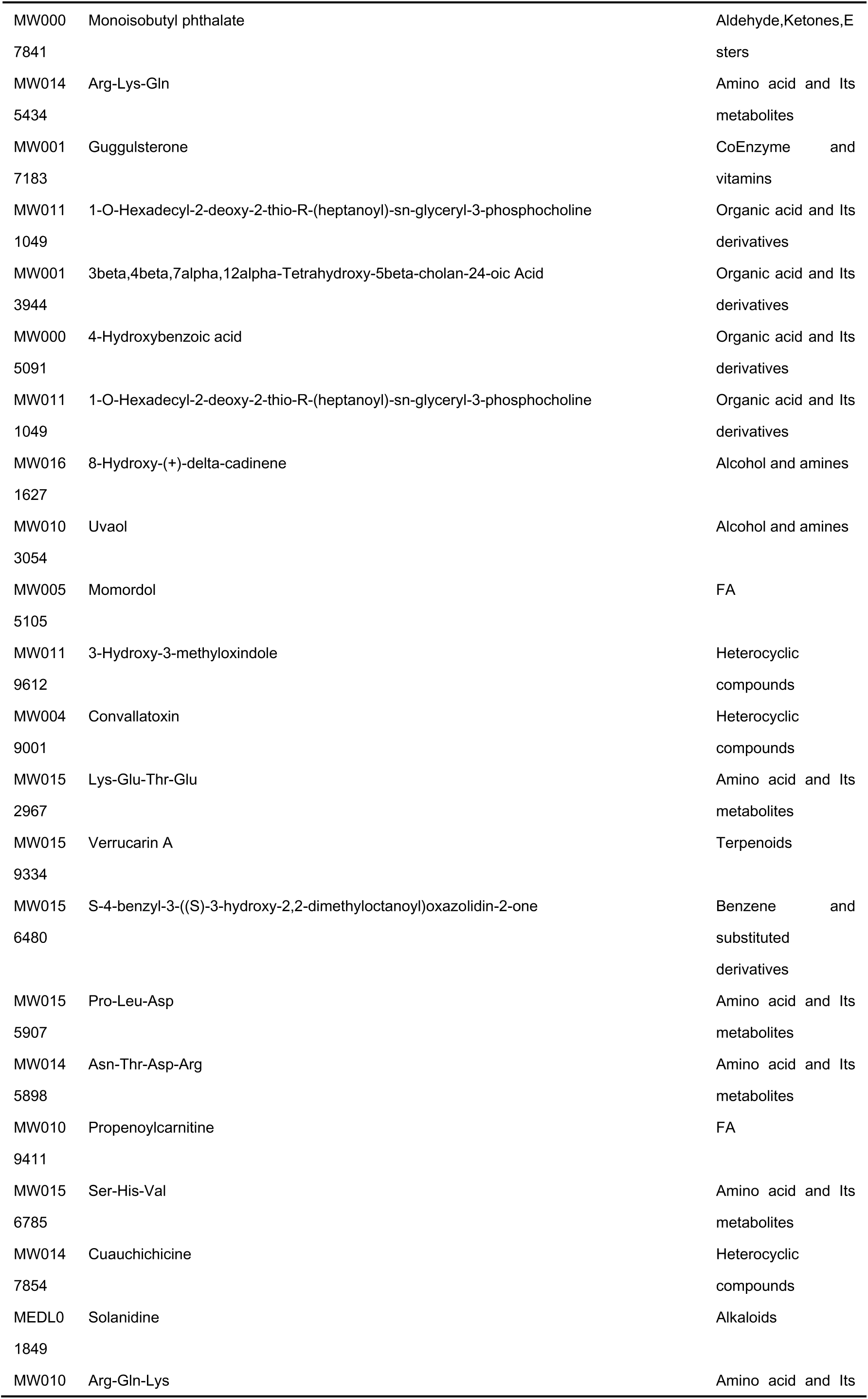
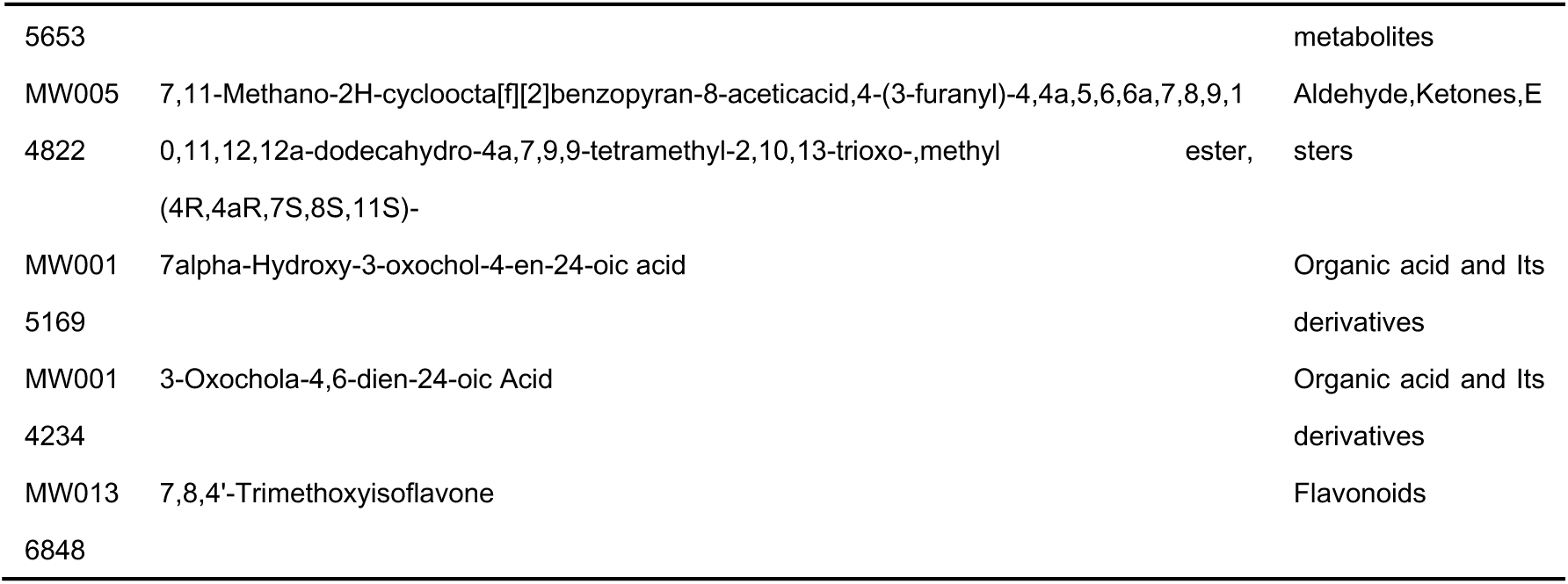
The information of top 10 metabolites with intergroup difference.

### Gut bacteria and metabolites

Gut bacterial communities exhibited divergent succession patterns across different T2D stages. In early healthy life stages, *s Bifidobacterium_pseudolongum*, *s Lactobacillus_johnsonii*, and *s Lactobacillus_reuteri* displayed higher abundances, which decreased progressively with age, in healthy/normal glucose stages. Notably, these three species exhibited proliferation in the T2D state, with their abundances further elevated following high-fat diet (HFD) feeding. We hypothesized that this phenomenon might be attributed to their involvement in regulating glycometabolism during both vigorous metabolic phases and disease states, as they showed positive correlations with specific BAs and amino acids.

Species from unclassified genera primarily proliferated in early healthy stages, including *g unclassified_Flavobacteriaceae*; *s gut_metagenome*, *g unclassified_Muribaculaceae*; *s mouse_gut_metagenome*, *g unclassified_Ruminococcaceae*; *s gut_metagenome*, *g unclassified_Lachnospiraceae*; *s Lachnospiraceae_bacterium_A2,* and *g unclassified_Clostridiaceae*; *s Candidatus_Arthromitus_sp*_SFB_rat_Yit. These taxa showed no association with BA or most amino acid metabolisms. A subset of species, such as *E. coli*, only proliferated in the death stage and exhibited positive correlations with certain BAs and amino acids.

Notably, species that increased exclusively in intervention groups (bT3I, RFI, MI) displayed uniform negative correlations with BAs and amino acids, including *g unclassified_Muribaculaceae*; *s gut_metagenome*, *g unclassified_Clostridia*; *s metagenome*, and *g unclassified_Christensenellaceae*; *s bacterium*_YE57. Collectively, these observations support a pattern: gut species that expand during unhealthy stages or in the deceased exhibit positive correlations with BAs and amino acids, whereas those enriched in healthy/normal glucose states show negative associations with these metabolites (Figure 7). For instance, *B. acidifaciens* positively correlated with gut BAs (e.g., glycoursodeoxycholic acid, sohyodeoxycholic acid, beta-muricholic acid, allocholic acid) and several amino acids, suggesting that BA metabolic pathways may play a critical role in glycometabolism regulation. Detailed abundance data for distinct gut species across the 9 groups are provided in Supplementary Table 5.

**Figure 7.**
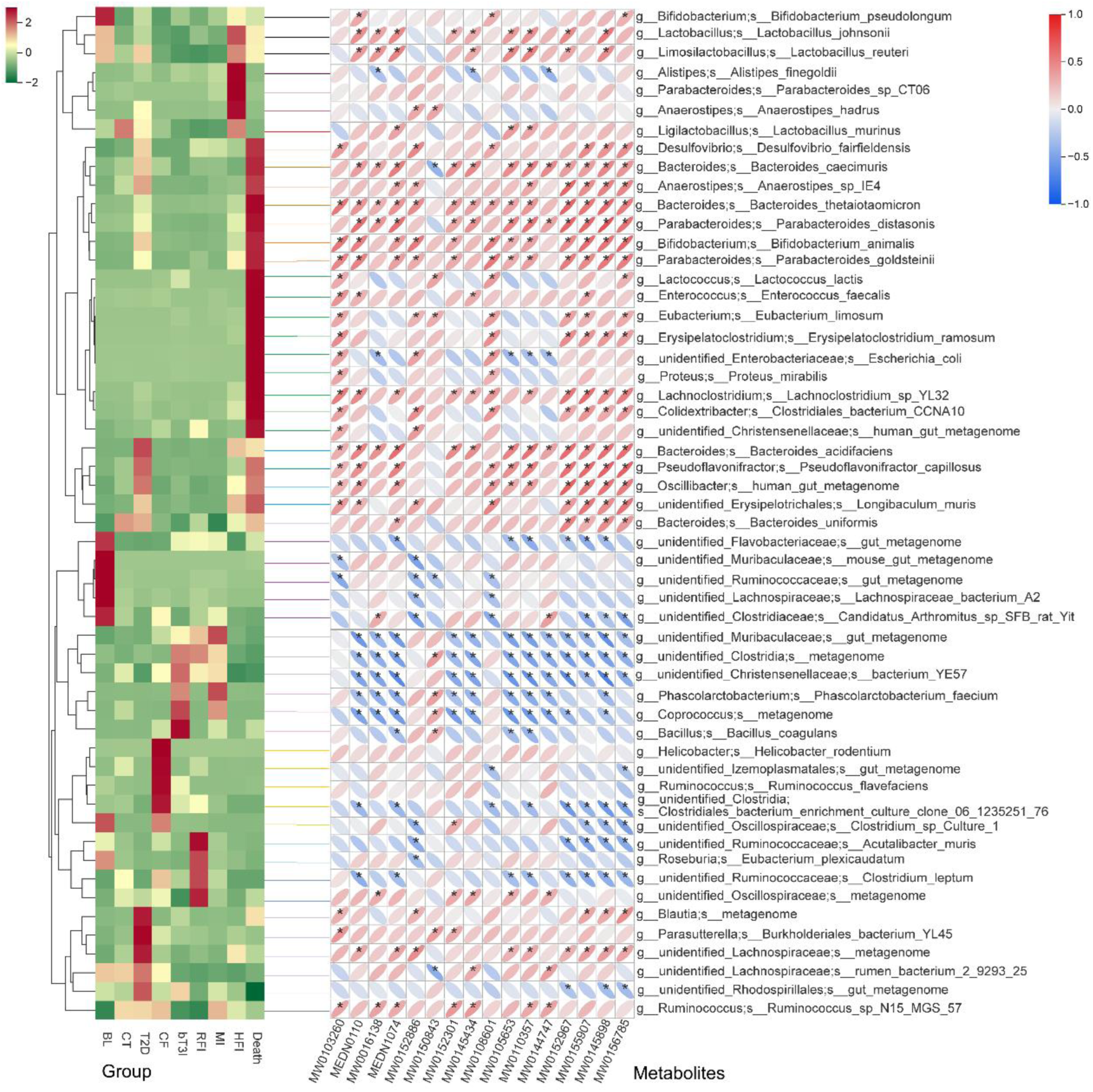
The results of correlation between the gut bacterial species with intergroup-difference and BAs (4), amino acids (12) which also had significant difference between groups, left heatmap is the relative abundance of the species and right part is correlation figure.

## Discussion

The results of the gut microbial diversity show that the older the more diverse, the reasons may be the gut microbiota relates to age and hormonal readiness, that has the same trend to human (21).To uncover the key metabolic mechanism of some specific gut microorgnisms in glycometabolism progress to develop probiotics using T2D prevention and therapy, we take more attention to the intergroup discrepancy of species levels. Actually, differences exist in the other levels and the rules of internal microbiota are comprehensive and complicated. We found the number of the *B. acidifaciens* had increased dramatically in the T2D stage. A study proved that the *B. acidifaciens* had an improved function to the colitis of mice as a probiotic (22). We speculate the reason probably is the T2D station promotes the occurring of colitis, and the species compensatory augmentation to protect the host because the host has a mechanism of self-protection. A new study demonstrated that *B. acidifaciens* treatment alone ameliorates liver injury through a bile salt hydrolase (23).

The number of glycoursodeoxycholic acid (GUDCA), a kind of secondary BAs, in T2D gut was increase, this result is consistent with other research (24), and it can improve various metabolic endpoints in mice with obesity. In fact, there were several BAs increasing in quantity in the T2D rats, like isohyodeoxycholic acid (isoHDCA, secondary BAs), beta-muricholic acid (β-MCA, primary BAs) and allocholic acid (ACA, primary BAs). However, these BAs showed different concentration in between T2D, HFI and Death three groups, GUDCA, β-MCA and ACA three BAs were higher in T2D and HFI than that in Death, but isoHDCA showed inverse trend comparing with the before ones. A research proved that the β-MCA can improve gut barrier function of male *Cyp2c70* knockout mice (25). Furthermore, a new intervention study showed that dietary supplementation with BA including ACA improves the health of shrimp and promotes growth and development by increasing the activity of intestinal digestive enzymes, hepatopancreas antioxidant enzymes, mRNA expression levels of gut-related immune genes and remolding gut microbiota (26). isoHDCA from feces is positive correlation with total cholesterol and high-density lipoprotein cholesterol of human (27). In a word, BAs perform a complicated functions involving combined action of liver and gut microbiota, some play protective roles to maintain the basic functions of hosts and the metabolites will be compensatory rise in quantity at the disease situations, like GUDCA, β-MCA and ACA.

Additionally, no matter which way to intervene the T2D rats, single fodder, β-T3 or MET, all could promote some specific gut microbial species increasing in number, but these bacteria do not have statistic correlation with the different BAs and amino acids, we conjectured these would regulate sugar metabolism by other metabolic pathways.

We designed a multi-group analysis and multi-omics testing to precisely uncover the rules of succession in the gut microbiota and the change of its metabolism in the whole progress of T2D. However, the results just can contribute some clues to find the key gut bacterial species and metabolic pathways, and it need more researches to prove that and come true using in human. Meanwhile, we did not test SCFAs because of the limitation of method, consequently it does not have the ability to describe the change rules of all gut metabolites.

### Conclusions

The succession pattern of gut microbiota exhibited divergent trends: diversity increased with age in healthy rats, whereas it decreased progressively in T2D rats. Notably, healthy diets, MET, and β-T3 interventions could restore the diminished microbial diversity in T2D rats. Concurrently, the composition of dominant gut bacteria also displayed stage-specific variations across different health phases. Among these, *B. acidifaciens* displayed the highest relative abundance specifically in the T2D state, suggesting its potential as a biomarker for T2D. Gut metabolic profiles were distinct across groups, with the T2D group showing an opposite metabolic signature compared to healthy controls. Importantly, healthy diets, MET, and β-T3 interventions partially reversed these aberrant metabolic features in T2D rats. *B. acidifaciens* exhibited positive correlations with four specific Bas, glycoursodeoxycholic acid, sohyodeoxycholic acid, beta-muricholic acid, and allocholic acid, mplying that the BA metabolic pathways associated with this bacterium may represent a critical regulatory mechanism underlying T2D pathophysiology.

## Abbreviations

T2D: type 2 diabetes mellitus
β-T3: β-tocotrienol, a kind of vitamin E
MET: metformin hydrochloride
FPG: Fasting blood glucose
BAs: Bile acids
SCFA: Short-chain fatty acids
FXR: Farnesol X receptor
LPS: Lipopolysaccharide
MF: Maintenance fodder
TMF: T2D model fodder
BL: Base line group, healthy rats
CT: Control group, feeding healthy rats by maintenance fodder for 17 weeks
CF: Control group, feeding healthy rats by maintenance fodder for 21 weeks
RFI: Intervention group, feeding T2D rats by maintenance fodder instead of T2D model fodder
bT3I: Intervention group, feeding T2D rats by maintenance fodder and β-tocotrienol (0.25mg dissolving in 1ml corn oil and feeding that to each rat by gavage every day) instead of T2D model fodder
MI: Intervention group, feeding T2D rats by maintenance fodder and metformin hydrochloride (45mg each mouse each day) instead of T2D model fodder
HFI: T2D rats which were continually fed by T2D model fodder
CTAB: Cetyltrimethylammonium Bromide
ASV: Amplicon sequence variants
NMDS: Non-Metric Multi-Dimensional Scaling
KEGG: Kyoto Encyclopedia of Genes and Genomes
PICRUSt2: Phylogenetic Investigation of Communities by Reconstruction of Unobserved States
LC/MS: Liquid Chromatograph/Mass Spectrometer
HPLC: High Performance Liquid Chromatography
PCA: Principal Component Analysis
HCA: Hierarchical Cluster Analysis
ANOVA: Analysis of Variance
PCoA: Principal Coordinates Analysis
GUDCA: Glycoursodeoxycholic acid
IsoHDCA: Isohyodeoxycholic acid
β-MCA: Beta-muricholic acid
ACA: Allocholic acid

## Acknowledgements

We express our appreciation to Wuhan Metware Biotechnology Co., Ltd for their invaluable assistance in sample detection.

## Authors’ contributions

Yanchao Liu and Han Bao performed the bioinformatics analysis and drafted the manuscript; Yanchao Liu and Jinni Yao performed the animal experiments. Yang Jiao, Hailing Li and Tao Yan performed data management and analysis. All authors reviewed the final manuscript.

## Data availability

The FPG and weight measurements dataset and Differential Metabolites Dataset are available in Supplemental table 1 and 6; The Functional microbial gene prediction dataset are available in Supplemental table 2-4; The Differential Gut Microbial Species Dataset are available in Supplemental table 5; The 16S rDNA sequencing data are available in China National Center (CNCB – Home) for Bioinformation via CRA025677.

## Declarations

### Ethics approval and consent to participate

The study protocol received approval from the Ethics Committee of Inner Mongolia Medical University (reference number: YKD202403024, signed on 20/07/2024). The number of experimental rats satisfied minimum sample size requirements. The present study was conducted in compliance with the Code of Ethics of the World Medical Association (Declaration of Helsinki) for Animal Experimentation.

## Consent for publication

Not applicable.

